# Histone deacetylase 9 promoter hypomethylation associated with adipocyte dysfunction is a statin-related metabolic effect

**DOI:** 10.1101/849695

**Authors:** Amna Khamis, Raphael Boutry, Mickaël Canouil, Sumi Mathew, Stephane Lobbens, Hutokshi Crouch, Toby Andrew, Amar Abderrahmani, Filippo Tamanini, Philippe Froguel

## Abstract

**Background:** Adipogenesis, the process whereby preadipocytes differentiate into mature adipocytes, is crucial for maintaining metabolic homeostasis. Cholesterol lowering statins increase type 2 diabetes (T2D) risk possibly by affecting adipogenesis and insulin resistance but the (epi)genetic mechanisms involved are unknown. Here, we characterised the effects of statin treatment on adipocyte differentiation using *in vitro* human preadipocytes cell model to identify putative effective genes.

**Results:** Statin treatment during adipocyte differentiation caused a reduction in key genes involved in adipogenesis, such as *ADIPOQ, GLUT4* and *ABCG1*. Using Illumina’s Infinium ‘850K’ Methylation EPIC array, we found a significant hypomethylation of cg14566882, located in the promoter of the histone deacetylase 9 (*HDAC9*) gene, in response to two types of statins (atorvastatin and mevastatin), which correlates with an increased *HDAC9* mRNA expression. HDAC9 is a transcriptional repressor of the cholesterol efflux *ABCG1* gene expression, which is epigenetically modified in obesity and prediabetic states. Thus, we assessed the putative impact of *ABCG1* knockdown in mimicking the effect of statin in adipogenesis. *ABCG1* KD reduced the expression of key genes involved in adipocyte differentiation and decreased insulin signalling and glucose uptake. In human blood cells from two cohorts, *ABCG1* expression was impaired in response to statins, confirming that *ABCG1* is in vivo targeted by these drugs.

**Conclusions:** We identified an epigenetic link between adipogenesis and adipose tissue insulin resistance in the context of T2D risk associated with statin use, which has important implications as HDAC9 and ABCG1 are considered potential therapeutic targets for obesity and metabolic diseases.

## 1 Background

Adipose tissue plays a crucial role in regulating insulin sensitivity and glucose homeostasis (1). In obesity, adipose expansion occurs as a result of cellular hypertrophy, *i.e.*, the increase in size of the adipocyte, and/or *de novo* adipogenesis, which is the production of new mature adipocytes from residing preadipocytes (2–4). Dysregulation in the adipogenic process is associated with metabolic diseases and insulin resistance (5) and is an independent risk factor for type 2 diabetes (T2D) (6). In contrast, appropriate adipocyte expansion is protective against T2D in the context of obesity (7,8). Adipogenesis occurs as a result of metabolic cues that trigger the induction of key differentiation regulators, such as ADIPOQ, FASN, PPARg, ABCG1 and GLUT4 (9–12). Epigenome wide association studies (EWAS) have found that hypermethylation within one of these genes involved in adipogenesis, *ABCG1*, was associated with increased body mass index (BMI), insulin resistance and T2D risk (13–16), opening avenues in the elucidation of the links between adipogenesis and metabolic diseases.

One of the most common drugs known to modulate adipogenesis are statins (17). The clinical use of statin is associated with an increased risk of insulin resistance and T2D risk (18), but the molecular mechanisms involved remain poorly understood. We hypothesised that statin treatment modulates adipogenesis by modifying the adipocyte epigenome. In this study, we confirmed the inhibitory effects of statin treatment in human preadipocytes and investigated the whole methylome to identify potential regulators that may be involved in adipogenesis.

## 2 Results

### 2.1 Statin treatment reduced adipogenesis and insulin signalling

The Simpson-Golabi-Behmel syndrome (SGBS) human preadipocyte cell line was used in this study as an *in vitro* model for adipocyte differentiation. In SGBS cells, lipid droplet formation occurred by 12-14 days of differentiation together with an increase in the expression of key adipogenic markers (19). We retrieved adequate SGBS cell morphology modification and formation of lipids droplets by day 12 (Additional File 1: Figure S1a), and observed that the expression of key genes involved in adipocyte differentiation and maturation was accordingly up-regulated (Additional File 1: Figure S1b).

For statin treatment, SGBS cells were differentiated for 6 days and then treated with atorvastatin and mevastatin for an additional 6 days until final maturation (Figure 1a). We observed a clear decrease in lipid-droplet formation in statin-treated SGBS cells (both atorvastatin and mevastatin) when compared to DMSO-vehicle controls (Figure 1b). We also found that statin-treatment induced a significant down-regulation of many key genes associated with adipogenesis reported above (*ABCG1, LEPTIN* and *GLUT4*), with the particular exclusion of *PPARG*, a gene known to play a role only in the early stages of adipocytes differentiation (Figure 1c). We then measured the effects of statin on downstream regulators of insulin signalling and found decreased efficiency of insulin to activate ERK and AKT (Figure. 1d). Taken together, the data support the inhibitory effects of statin in the human adipocyte differentiation and insulin signalling, a similar effect reported in statin-treated 3T3-L1 mouse adipocyte cells(17).

**Figure 1:**
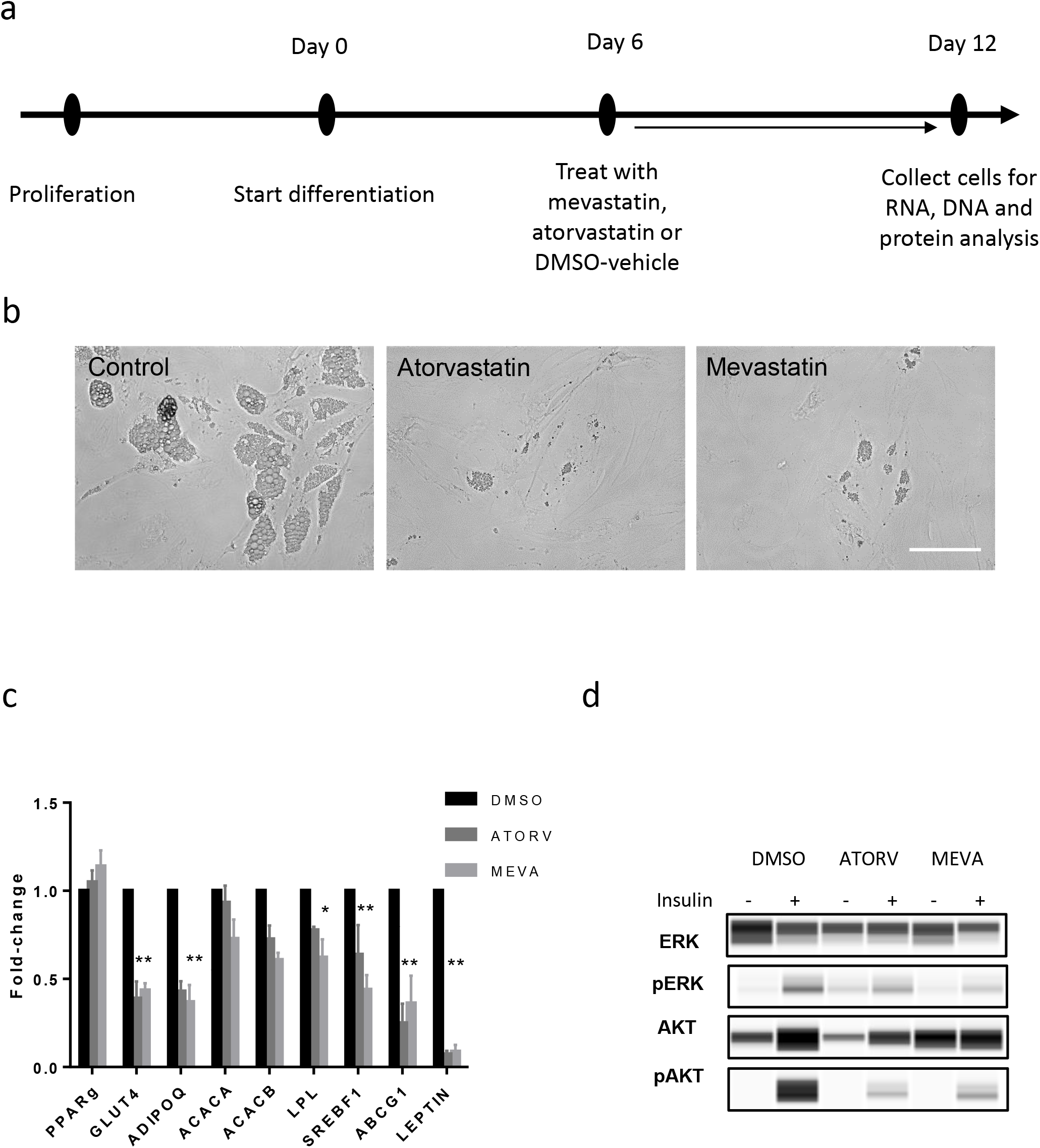
Response of SGBS cell line to statin treatment. (a) The method used in treating the SGBS cell line at day 6 of differentiation for 6 days. (b) Microscope images showing lipid droplets in statin-treated cells at day 12 of differentiation following statin treatment for atorvastatin, mevastatin and DMSO controls (×10 magnification scale bar 10 μm). (c) Expression of key adipose genes for statin-treated cells compared to time-matched DMSO controls (normalised to housekeeping gene B2M). * p < 0.05; ** p < 0.01 (d) Protein expression of insulin signalling proteins pAkt and pErk in statin-treated cells compared to controls using WES.

### 2.2 Whole methylome analysis of statin-treated SGBS cell line

To identify potential regulators involved in statin-induced adipocyte dysregulation, we performed an unbiased whole methylation analysis in statin-treated SGBS cells using Illumina’s Infinium ‘850K’ Methylation EPIC arrays (Additional File 1: Figure S2). We filtered differentially methylated positions (DMPs) located in the promoter region, annotated as TSS200 or TSS1500 (within 200-1500 base pairs from the transcription start site), in order to identify DMPs that were likely to have a biological effect. The most significant DMP was cg14566882, located in the promoter of the histone deacetylase (*HDAC9*) gene, in mevastatin-treated cells (β = 8.28 %; p = 5.55 × 10^−6^) (Figure 2a). This DMP was also significantly hypomethylated in response to atorvastatin treatment, compared to DMSO-vehicle controls (β = 5.53 %; p = 1.35 × 10^−3^) (Figure 2a, b).

**Figure 2:**
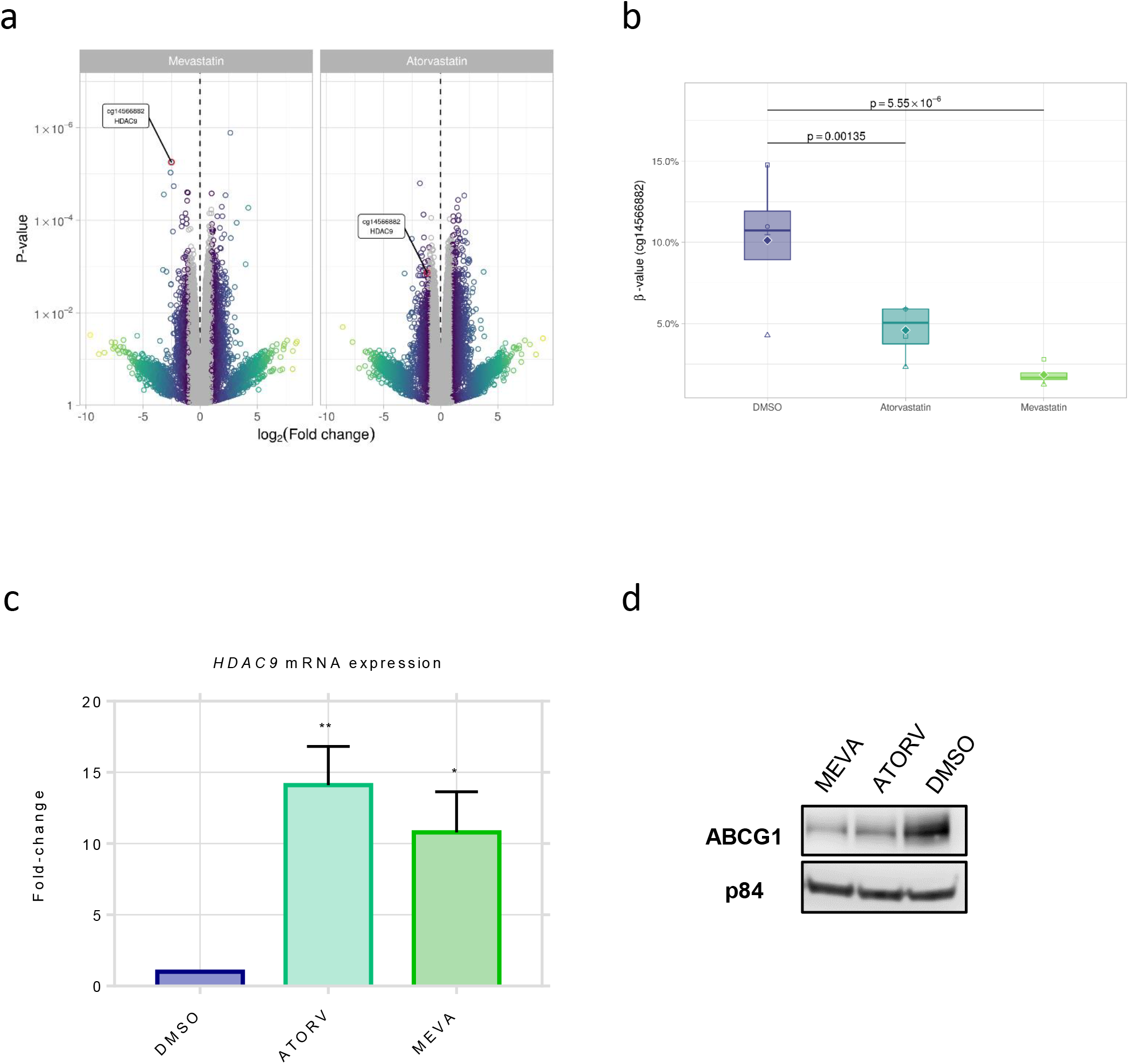
Whole methylome analysis of statin-treated SGBS cells. (a) Volcano plots of whole methylome results for statin-treated cells (grey indicates log2 fold change < 1). (b) The hypomethylation of the cg14566882 CpG within the *HDAC9* gene in atorvastatin and mevastatin-treated cells compared to vehicle-treated DMSO cells in the 4 biological replicates (raw β-values shown). (c) The mRNA expression level of *HDAC9* in mevastatin and atorvastatin-treated SGSB cell line. * p < 0.05; ** p < 0.01 (d) The protein expression of ABCG1 compared to housekeeping gene p84 shows a reduced expression in atorvastatin and mevastatin-treated SGBS cells.

Additionally, 87 DMPs were shared between the mevastatin and atorvastatin-treated groups (Additional File 2: Table 1) and of the hypomethylated DMPs, cg14566882 in *HDAC9* remained the top candidate in both atorvastatin and mevastatin treatments (Additional File 1: Figure S3). A significant differentially methylated region (DMR) overlapping this promoter region was also found in response to both treatments (Additional File 1: Figure S4; False discovery rate < 0.05). In order to validate the effect of cg14566882 hypomethylation on the expression of the *HDAC9* gene, we performed qPCRs in statin-treated SGBS cell lines and found significant up-regulation of the *HDAC9* gene at the mRNA level (p < 0.05; atorvastatin: 14-fold; mevastatin:11-fold) (Figure 2c).

ABCG1 has been reported to be regulated by HDAC9-mediated changes in acetylation (20,21) and may be targeted by the *HDAC9* epigenetic alteration. We confirmed that ABCG1 protein expression is indeed down-regulated in response to mevastatin and atorvastatin treatment (Figure 2d). We confirmed that in response to statin, this effect was independent of the previously reported hypermethylation in cg06500161 (p-value > 0.5) and cg27243685 (p-value > 0.5) found to be associated with increased BMI and T2D incidence (Additional File 1: Table 2) (15,22–24).

### 2.3 Knockdown of *ABCG1* in SGBS preadipocytes reduced adipocyte differentiation

We performed an *ABCG1* knockdown (KD) in SGBS cells to address whether reduced *ABCG1* expression mimics the effect of statins in adipogenesis. SGBS preadipocytes were stably transfected with a shRNA targeting *ABCG1* mRNA and followed them quantitatively through maturation and differentiation and this data was compared with cells transfected with a non-targeting shRNA (control). A similar protocol for stably knockdown *Abcg1* via shRNA has previously been achieved and described in mouse 3T3-L1 preadipocytes (25). We initially analysed the expression of ABCG1 protein in normal adipocytes to show that it is positively associated with adipogenesis as ABCG1 starts to become expressed at day 6 of differentiation (Figure 3a). The efficient silencing of *ABCG1 (ABCG1* KD) was confirmed at the protein level (Figure 3b). *ABCG1* KD in SGBS cells was accompanied by a significant reduction in the lipid content (20 % reduction, p<0.05; Figure 3c), along with the down-regulation of the following adipocyte differentiation markers *FASN, PPARG* and *PLIN1* and key adipocyte maturation markers *ADIPOQ* and *GLUT4* (Figure 3d). As a consequence of impaired adipogenesis, the *ABCG1* KD led to a significant reduction in glucose uptake stimulated by insulin (65 % reduction, p < 0.001; Figure 3e). In addition, we found a decreased efficiency of insulin to activate AKT in those cells (Figure 3f). As a whole, this data indicates that *BCG1* levels are pivotal for the control of human adipocyte differentiation and glucose metabolism.

**Figure 3:**
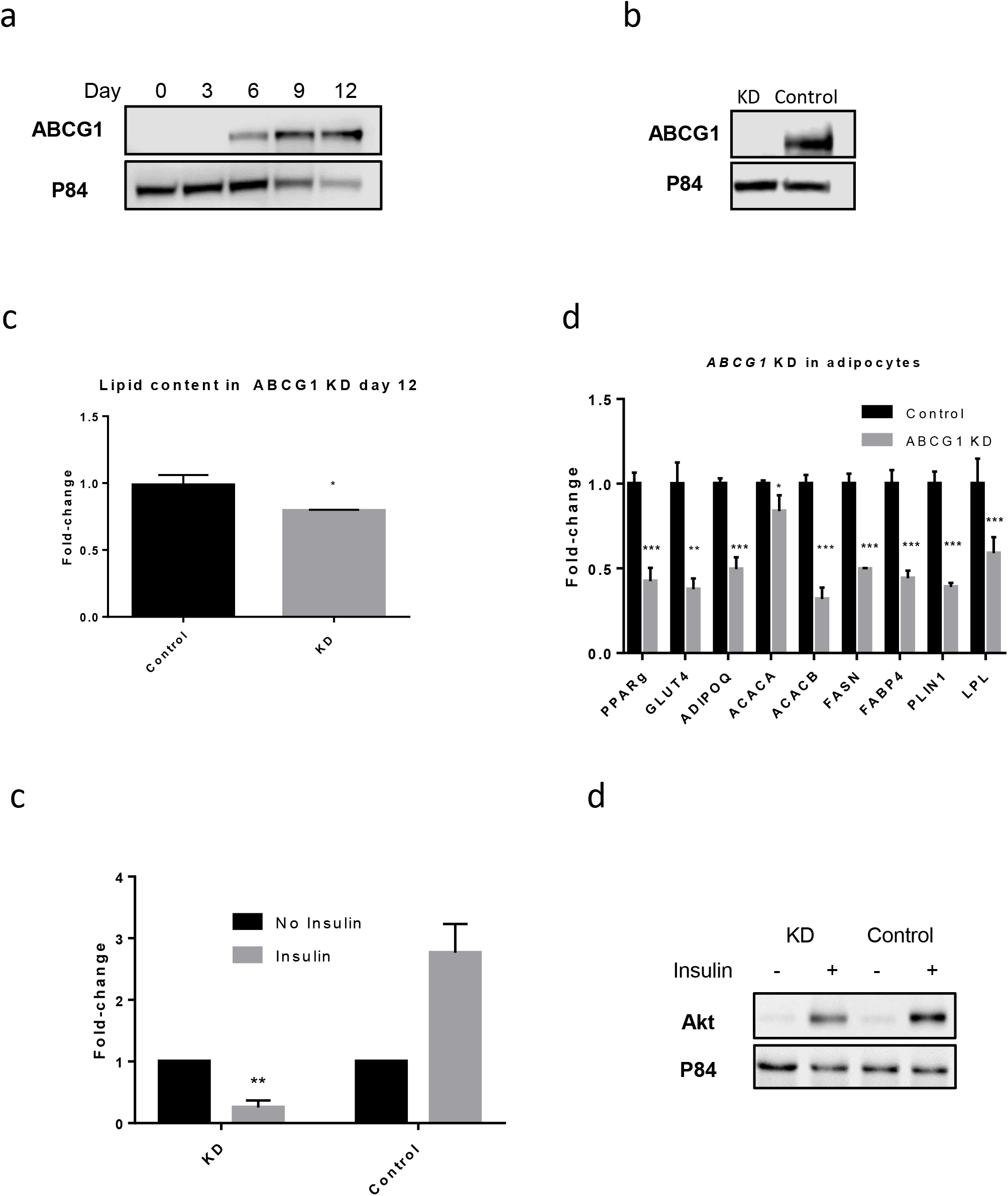
Adipogenesis changes in stably transfected *ABCG1* KD cells. (a) Western Blot protein expression of ABCG1 (a) during differentiation (b) after silencing in SGBS cell lines. (c) Lipid content analysed by red oil of *ABCG1* KD compared to controls. (d) Expression of key adipose genes at day 12 differentiation in KD *ABCG1* cells compared to shRNA controls, normalised to housekeeping gene *B2M* and compared to DMSO-vehicle controls. Experiments were performed at n = 4 biological replicates. (e) Glucose uptake in *ABCG1* KD compared to controls stimulated with or without 1 μM insulin for 1 hour. Fold change in KD and control cells compared to cells not treated with insulin. (f) Analysis of insulin signalling in SGBS *ABCG1* KD cell line through protein expression of phosphorylated AKT at day 12, stimulated with or without 200 nM insulin for 1 hour, using western blot analysis. * p < 0.05; ** p < 0.001; *** p < 0.0001.

### 2.4 *ABCG1* is down-regulated in response to statin in human blood samples

We next explored whether *ABCG1* was also dyregulated in samples from human subjects. We analyzed reported transcriptomic data from blood samples from two cohorts. The first consisted of a total of 57 individuals from the ECLIPSE cohort, of which 13 were statin users (26). A significant reduction in the expression of *ABCG1* in the statin group (p = 1.41 × 10^−5^) was found, compared to non-users (Figure 4a). In addition, we also analysed data from the YELLOW II study (27), a retrospective study following 85 individuals before and following an extensive 8-12 week statin therapy. In peripheral blood mononuclear cells obtained from blood samples, *ABCG1* expression was significantly decreased following statin treatment, compared to baseline levels, for two *ABCG1* probes (ILMN_1794782 p = 2.76 × 10^−5^; ILMN_2329927 p = 4.28 × 10^−4^) (Figure 4b). Taken together, this demonstrates that *ABCG1* reduction in response to statin is indeed reflected in human blood samples. Of note, no data on *HDAC9* expression was available in the ECLIPSE case control study, and no significant change in *HDAC9* expression was reported in the intervention YELLOW II study, maybe due to the lack of sufficient statistical power of this study.

**Figure 4:**
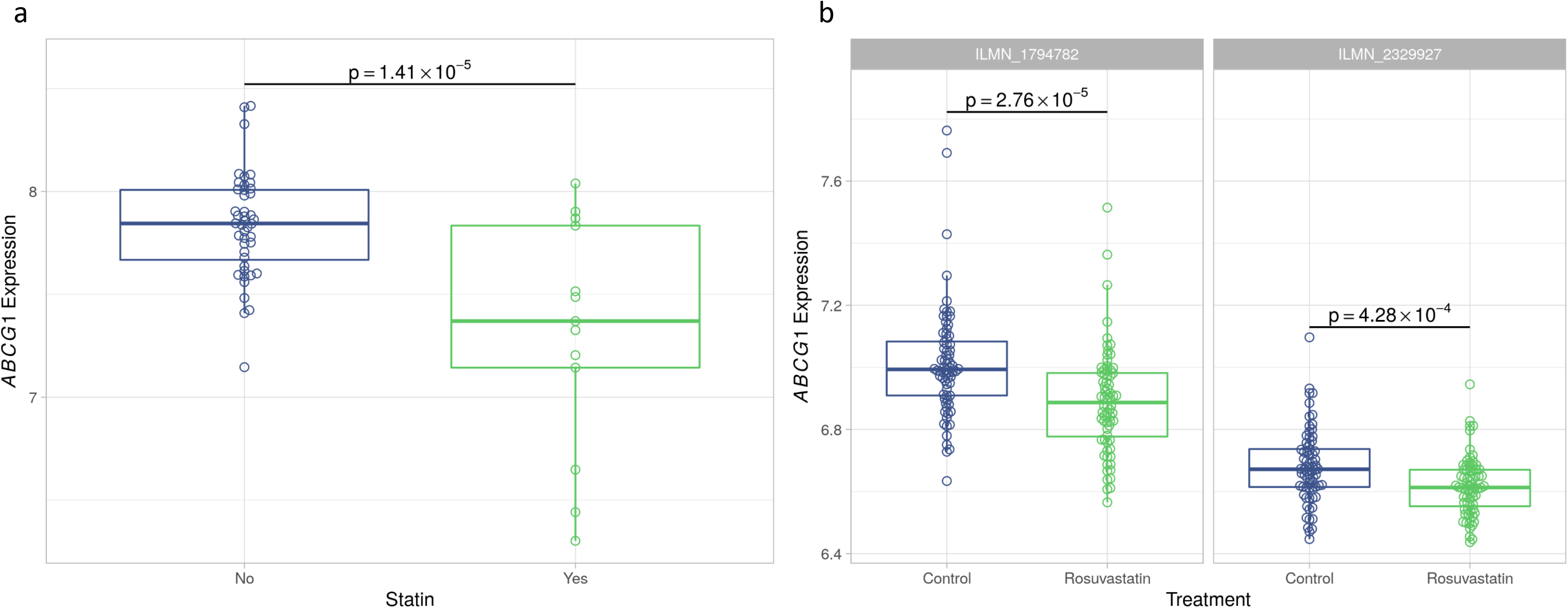
The expression of *ABCG1* in human samples. (a) *ABCG1* expression was reduced in 13 statin-treated individuals compared to control non-users using transcriptomic data. (b) Data from a total of 85 samples, after extensive statin treatment for 8-12 weeks, there was a reduction in *ABCG1* expression compared to baseline levels in two probes.

## 3 Discussion

A recent 15 year prospective study found a staggering 38 % increased incidence of T2D in statin users, regardless of the type of statin used (28). Here, we report that two statins, atorvastatin and mevastatin, hamper the differentiation process in the SGBS human preadipocyte cell line and decreased insulin sensitivity.

We focused our analysis on promoter DMPs, which are normally inversely correlated with expression (29,30). Therefore, not surprisingly, given the inhibitory effect of statin, our whole methylome analysis revealed that most DMPs were hypermethylated. This includes the *IDI1* gene, which encodes the isopentenyl diphosphate isomerase, a component of the cholesterol synthesising pathway (31,32).

We report for the first time that statin treatment was associated with a significant hypomethylation of *HDAC9* promoter, which is inversely correlated with *HDAC9* gene expression. These findings are of particular significance in light of several studies that demonstrated the key role of HDAC9 in adipocytes function: overexpression of *Hdac9* in 3T3-L1 preadipocyte mouse cell lines suppressed adipogenesis and inversely, preadipocytes isolated from *Hdac9* knockout mice had an accelerated adipocyte differentiation (33). Furthermore, *Hdac9* knockout mice showed improved metabolic homeostasis and were protected from adipose tissue dysfunction in mice fed on a high fat feeding (34). These studies clearly indicate the deleterious role of HDAC9 in maintaining adipocytes homeostasis both *in vitro* and *in vivo*.

*HDAC9*-deficient macrophages and monocytes directly increased the accumulation of total acetylated H3 and H3K9 at the promoter of the *ABCG1* gene (20,21), thereby inducing the transcription of the *ABCG1* gene, indicating that HDAC9 mediates the expression of *ABCG1* through promoter-mediated acetylation. This is of particular interest, as several studies have reported a role of ABCG1 in obesity, insulin resistance and T2D. Elevated *ABCG1* expression is associated with increased fat mass from obese individuals, suggesting that *ABCG1* is also involved in human adipogenesis (25). Although genome-wide association studies have not found any single nucleotide polymorphisms (SNPs) within or nearby *ABCG1* associated with increased T2D risk, several EWAS have found that hypermethylation in the *ABCG1* gene was associated with fasting glucose, HbA1C levels, lipid metabolism, fasting insulin, T2D risk and BMI (15,16,22,35–38). Additional observations in mouse models have shown that Abcg1^−/−^ mice were protected from high fat diet-induced glucose intolerance (39). A recent study found that *ABCG1* expression is reduced in both subcutaneous and visceral adipose tissue in morbidly obese patients with metabolic syndrome compared to those without metabolic syndrome, providing further evidence for a role of *ABCG1* in the maintenance of metabolic homeostasis in adipocytes (40). In addition, two studies showed that *ABCG1* expression was decreased in blood white human cells in response to statins. As ABCG1 was down-regulated in response to statin, we hypothesised that ABCG1 plays a role in statin-induced adipocyte dysregulation.

Indeed, we showed that *ABCG1* expression increases during human SGBS adipocyte differentiation and through *ABCG1* silencing, confirm that the level of *ABCG1* expression is crucial for the appropriate expression of lipid metabolism markers, which include *FASN, FABP4, PLIN1* and *PPARG*, for correct human adipocyte differentiation. Our findings are consistent with previous data showing variation in these four genes following *Abcg1* silencing in mouse 3T3-L1 pre-adipocyte cells (41). The down-regulation of *GLUT4* in *ABCG1* KD suggested a decrease in insulin-induced glucose uptake. Indeed, we confirmed a down-regulation of phosphorylation of AKT and ERK. Collectively this data indicates that normal ABCG1 function is required for adipogenesis and insulin signalling. In addition, we have confirmed using two separate datasets that statin use was indeed correlated with a reduction in *ABCG1* expression in human blood samples. Other studies have reported a link between ABCG1 downregulation and diabetes incidence (42) and high fasting glycaemia (43). Taken together, our human cellular data is consistent with human observational studies, in which the inhibition of *ABCG1* expression was deleterious for metabolism in adipose tissue.

## Conclusions

The model proposed based on our data from statin-induced insulin resistance is hypomethylation of the *HDAC9* promoter induces *HDAC9* gene expression, which in turn blocks *ABCG1* expression and thereby adipocyte differentiation and metabolic dysfunction (Figure 5). Adipocyte turnover by adipogenesis is crucial for the maintenance of metabolic homeostasis and insulin sensitivity (44). Our data provides a novel epigenetic link between adipogenesis dysfunction and insulin resistance, mediated by statins. The increased understanding of adipogenesis provides a promising new avenue for the treatment of metabolic disease in obesity (9,44). Both HDAC9 and ABCG1 have been proposed as therapeutic targets for patients with obesity in separate previous studies (34,41), however, our data support a mechanistic pathway linking them to metabolic diseases.

**Figure 5:**
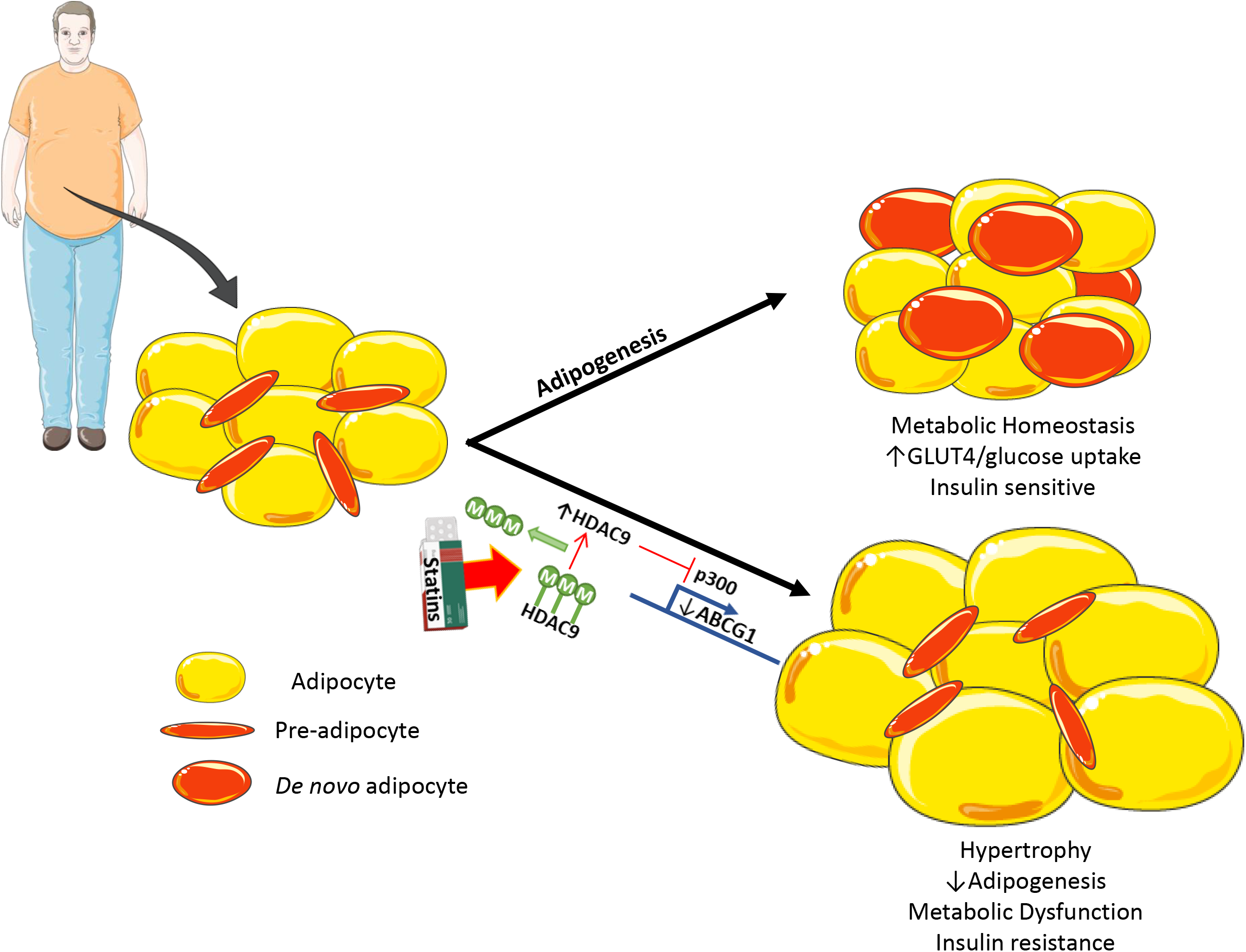
A schematic representation of the role of adipocyte turnover in health and disease. In healthy individuals, preadipocytes differentiate into mature adipocytes, which have a role in maintaining insulin sensitivity. However, in response to statins, epigenetic changes in *HDAC9* cause acetylation changes in *ABCG1* and other crucial adipogenesis genes, which in-turn an obstruction of differentiation and metabolic dysfunction.

## 4 Methods

### 4.1 Cell culture and differentiation of SGBS cell line

SGBS human preadipocyte cell line was kindly provided by Prof. Dr. M. Wabitsch (University of Ulm, Germany) and maintained in DMEM/F12 supplemented with 10 % foetal bovine serum and 0.01 % penicillin/streptomycin (15140-122 Life Technologies), as previously described (19). Confluent preadipocytes were differentiated under serum-free culture conditions by washing twice with phosphate buffered saline (PBS) and then exposing to DMEM/F12 supplemented with 2 μmol/l rosiglitazone, 25 nmol/l dexamethasone, 0.5 mmol/l methylisobuthylxantine, 0.1 μmol/l cortisol, 0.01 mg/ml transferrin, 0.2 nmol/l triiodotyronin, and 20 nmol/l human insulin for 4 days. The cells were then cultured for a further 8 days in fresh DMEM/F12 supplemented with 0.1 μmol/l cortisol, 0.01 mg/ml transferrin, 0.2 nmol/l triiodotyronin, and 20 nmol/l human insulin. Microscopic images of SGBS cells were taken under a microscope (IT404; VWR) using the Motic Image plus version 2.0 (Motic Europe).

### 4.2 Treatment with statins

At 6 days of differentiation, SGBS cells were treated with 10 μM mevastatin (M2537, Sigma Aldrich) or atorvastatin (PZ0001, Sigma Alrich) and compared to a Dimethyl sulfoxide (DMSO)-vehicle control (D2650, Sigma Aldrich). The cells were then incubated for a further 6 days. On day 12 of differentiation, cells collected for further analysis.

### 4.3 Whole methylome analysis

DNA was extracted from SGBS cells treated with atorvastatin and mevastatin at day 6 for 6 days and collected the cells at day 12. DNA was extracted using the NucleoSpin Tissue kit (Takara Bio). Bisulfite conversion of 500 ng genomic DNA was performed using the EZ-96 DNA Methylation kit (Zymo Research) following the manufacturer’s protocol. Bisulfite-converted DNA was subjected to genome-wide DNA methylation analysis using Illumina’s Infinium ‘850K’ Methylation EPIC array to identify differentially methylated positions (DMPs). The resulting DNA methylation IDAT files were imported using the *minfi* R package for further processing and quality control (45). The following CpG probes were excluded from further analysis: probes on sex chromosomes, cross-hybridising probes, non-cg probes and probes that lie near single nucleotide polymorphisms (SNPs). Probe-design biases and batch effects were normalised using R packages *ENmix* (46) and *SVA* (ComBat) (47), respectively. To identify DMPs, the R package *limma* was used (48). The model included treatment (atorvastatin, mevastatin or DMSO-vehicle) as a categorical variable and replicates / day of experiment as a covariate. Methylation levels denoted by beta-values, where 0 indicates 0 % methylation and 1 indicates 100 % methylation, were transformed to M-values (49). To identify differentially methylated regions (DMRs), the R package *DMRcate* was used (50). *DMRcate* ranks the differentially methylated regions across the genome using Gaussian kernel smoothing based on the DMPs.

### 4.4 RNA extraction, cDNA conversion and RT-PCR

Total RNA was extracted from cultured cells using a RiboPure RNA Purification Kit (AM1924; Invitrogen), according to the manufacturer’s instructions and quantified on a NanoDrop Spectrophotometer (Thermo Scientific). RNA was reverse transcribed using a High-Capacity RNA-to-cDNA Kit (4387406; Applied Biosystems), according to the manufacturer’s instructions. qPCRs were conducted on an Applied Biosystems 7900HT Fast Real-Time PCR System and quantitative expression levels were obtained using the SDS v2.3 Software (Applied Biosystems) using Taqman Gene Expression Assays (ThermoFisher Scientific). The following probes were used: *ABCG1* (Hs01555193_m1), *CEBPB* (Hs00270923_s1), *LPL* (Hs00173425_m1), *ACACA* (Hs01046047_m1) *FABP4* (Hs01086177_m1), *GLUT4* (Hs00168966_m1), *ACACB* (Hs00153715_m1), *PPARG* (Hs01115513_m1), *CEBPA* (Hs00269972_s1), *ADIPOQ* (Hs00605917_m1), *FASN* (Hs01005622_m1), *SREBF1* (Hs01088691_m1) and *PLIN1* (Hs00160173_m1) (Life Technologies). Each reaction was normalised to a *beta-2-microglobulin* (*B2M*) control (Hs00984230_m1; Applied Biosystems). For quantifications using SYBR-green, qPCRs were performed using (SsoAdvanced Universal SYBR Green Supermix, BioRad) using the BioRad CFX96 Real-Time PCR Detection System (Biorad). *HDAC9* gene primer sequences were forward: AGTGGCAGAGAGGAGAAGCA and reverse: CAGTTCTCCAGGCTCTGGTC. Atleast three biological replicates were performed and the data represents the means ±SEM. A two-tailed t-test was performed using GraphPad Prism (GraphPad software Inc., La Jolla, USA) and p-value < 0.05 was considered to be statistically significant.

### 4.5 Lipid quantification

Culture medium was removed and the cells were washed twice with PBS and then fixed with 10 % formalin for 30 min. Fixed cells were washed twice with water and incubated in 60 % isopropanol for 2 min. The alcohol was discarded and the cells were then incubated in an Oil Red O (Sigma) and water solution (3:2) for 5 min. Cells were rinsed four times with water and then 100 % isopropanol was added to extract the red oil. The absorbance was measured at 540 nm on a VERSAmax ELISA microplate reader (Molecular Devices) and analysed using the SoftMax Pro Software (Molecular Devices).

### 4.6 Western blotting

Cells were washed twice in ice-cold PBS and harvested in RIPA buffer (ThermoFisher Scientific) supplemented with protease inhibitors. Cell lysates were centrifuged at 20,000 × *g* for 20 min at 4°C and the total protein lysate was quantified using Bradford Reagent (B6916; Sigma). Then, 40 μg of total protein lysate was separated on a 4-12 % Bis-Tris Plus Gel (Life Technologies) and transferred to a nitrocellulose membrane using the iBlot2 Gel Transfer Device (Life Technologies). Membranes were blocked in 0.5 % non-fat dry milk and probed with anti-ABCG1 (1:1000; ab52617, abcam) and antinuclear matrix protein p84 (1:5,000; ab487, abcam) primary antibodies anti-rabbit IgG (1:20,000; Ab205718, abcam) and anti-mouse IgG (1:5,000; A4416, Sigma Aldrich) secondary antibodies for 1 h at room temperature. Membranes were exposed using Clarity Western ECL Substrate (Bio-Rad) and protein bands were detected on a LI-COR Imaging system with C-DiGit Image Studio 4.0 software (LI-COR Biosciences, Ltd., UK).

### 4.7 Phosphorylated AKT analysis

Differentiated cells cultured in DMEM/F12 medium were serum starved overnight and then washed with PBS and stimulated with or without 200 nM insulin for 1 h in DMEM/F12 (without glucose or serum) at 37°C in 5 % CO_2_. The total protein was harvested in RIPA buffer supplemented with protease and phosphatase inhibitors, as described above. The primary antibodies (all used at 1:1,000 dilution unless otherwise stated) used were anti pAKT (S473; Cell Signaling) and anti Akt (9272 - 1:5,000 dilution; Cell Signaling) and the secondary antibody used was goat pAb to Rb igG (Ab205718 - 1:20,000 dilution; Abcam). Protein expression studies were also performed by WES, an automated capillary-based size separation and nanoimmunoassay system (ProteinSimple, San Jose CA, USA a Bio-Techne Brand), according to manufacturer’s protocol, for analysis performed using the Compass for Simple Western software v.4.0. The wes was performed on SGBS samples (1:100) for anti pAKT (S473; Cell Signaling), anti Akt (9272 - 1:100 dilution; Cell Signaling) anti pERK (9102; Cell Signaling) and anti ERK (9101 - 1:100 dilution; Cell Signaling).

### 4.8 Glucose uptake assay

A Glucose Uptake-Glo Assay (J1342; Promega) was used to measure glucose uptake in differentiated cells (day 12), according to the manufacturer’s protocol. A total of 20,000 cells in 100 μl media were plated in each well of a 96-well white plate. Differentiated cells were cultured overnight in DMEM/F12 media with no serum. On the day of the assay, the media was replaced with 100 μl DMEM/F12 (without glucose or serum) supplemented with or without 1 μM insulin and incubated for 1 h at 37°C in 5 % CO_2_. Cells were washed with PBS and 50 μl 1 mM 2-Deoxy-D-glucose (2DG) was added to each well and incubated for 10 min at room temperature. Next, 25 μl Stop Buffer was added followed by 25 μl Neutralization Buffer per well. Finally, 100 μl 2DG6P Detection Reagent was added and incubated for 4.5 h at room temperature. The luminescence was recorded on a Mithras LB 940 luminometer (Berthold Technologies) and analysed using the MicroWin Software (Berthold Technologies).

### 4.9 *ABCG1* silencing using shRNA lentiviral vector

Undifferentiated SGBS cells were plated at 50 % confluence in 6-well plates and infected with commercial lentiviral particles targeting either human *ABCG1* (TRCN0000420907; Sigma) (TAGGAAGATGTAGGCAGATTG) or non-target controls (SHC202) (CCGGCAACAAGATGAAGAGCACCAACTC) and (TRCN0000158395; CCTACAGTGGATGTCCTACAT) (Sigma Aldrich). The transduced cells were selected in media containing 1 ug/ml puromycin for 6 days. Stable *ABCG1* KD and control cells were then cultured and differentiated into mature adipocytes, as described above.

### 4.10 Analysis of transcriptomic data from human samples

Transcriptomic data (Affymetrix Human Gene 1.1 ST Array) from the GSE71220 dataset was downloaded from the GEO database. The subjects analysed were the 57 control samples from the Evaluation of COPD Longitudinally to Identify Predictive Surrogate Endpoints (ECLIPSE) study, of which 13 were statin-users (26). In addition, transcriptomic data (Illumina HumanHT-12 WG-DASL V4.0 R2 expression beadchip) from peripheral blood mononuclear cells from the YELLOW II retrospective study (GSE86216) was also downloaded from the GEO dataset. This included blood samples from a total of 85 patients that were analysed before and after an extensive 8-12 week statin treatment (27). For both datasets, the data was downloaded and analysed using the R packages GEOquery (45) and limma (48).

## List of abbreviations

B2M: beta-2-microglobulin
BMI: body mass index
DMP: differentially methylated position
DMR: differentially methylated region
DMSO: dimethyl sulfoxide
EWAS: epigenome wide association study
HDAC9: histone deacetylase 9
KD: knockdown
SGBS: Simpson-Golabi-Behmel syndrome
SNP: single nucleotide polymorphism
T2D: type 2 diabetes

## Declarations

### Ethics approval and consent to participate

Not applicable

### Consent for publication

Not applicable

### Availability of data and materials

The datasets generated and/or analysed during the current study are available in the Gene Expression Omnibus (GEO) repository, under GSE139211 https://www.ncbi.nlm.nih.gov/geo/query/acc.cgi?acc=GSE139211. Access to datasets will remain private during period of manuscript review. Reviewers can access data using the following secure token: yzmzwwkaprgpdoj

### Authors’ contributions

AK, AA, TA, FT and PF designed the project. AK, FT, SM, RB and HC have performed the experiments. SL performed the methylome wet lab experiments and MC performed the methylation analysis. AK and MC prepared the figures. AK and PF wrote the manuscript. All authors edited the paper.

## Acknowledgements

The authors would like to thank Prof. Dr. M. Wabitsch (University of Ulm, Germany) for kindly providing the SGBS cell line and differentiation protocol.

