## Additional File 1 for "Histone deacetylase 9 promoter hypomethylation associated with adipocyte dysfunction is a statin-related metabolic effect"

### Slide 1
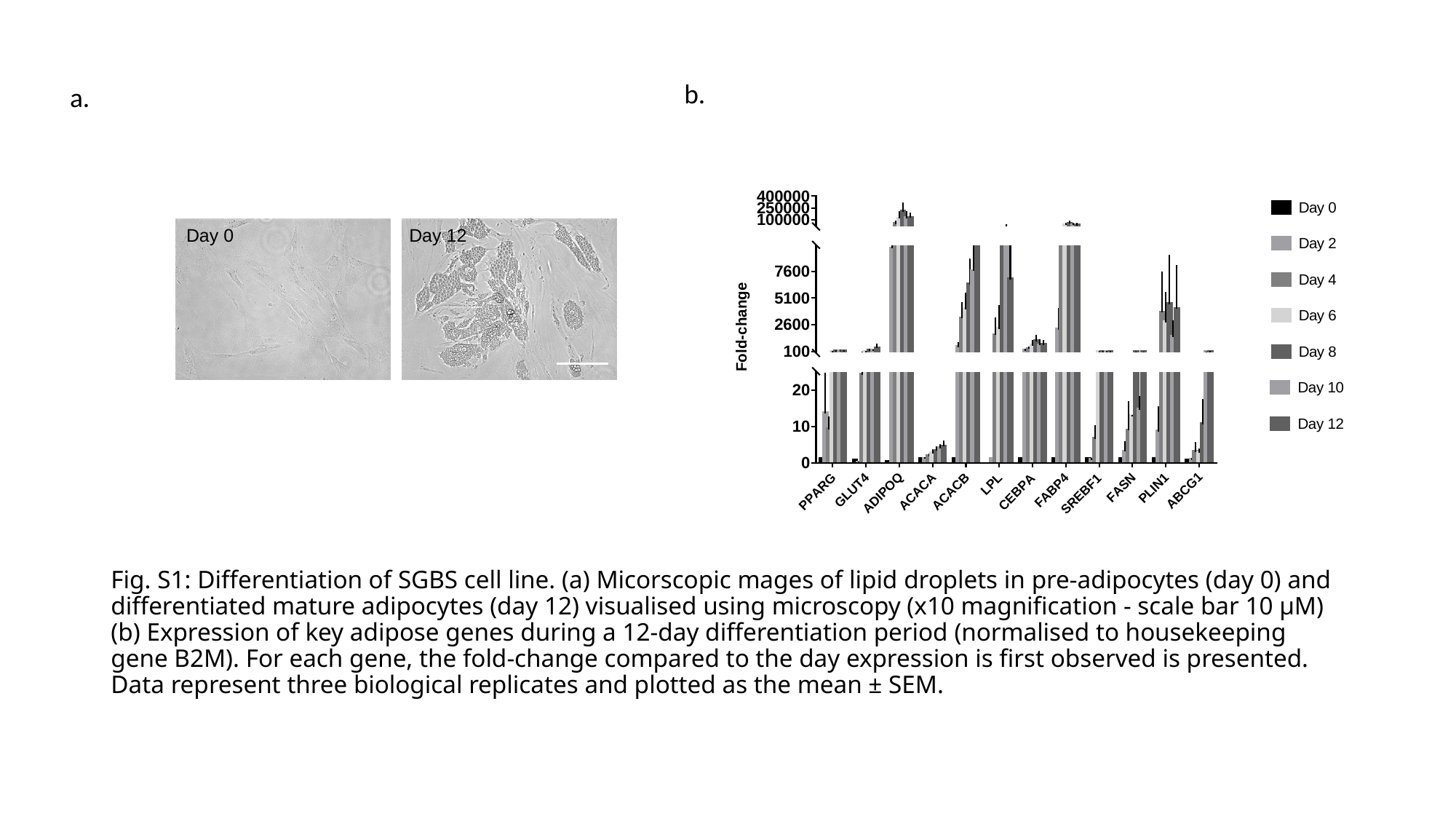

b.
a.
Day 0
Day 12
#### Fig. S1: Differentiation of SGBS cell line. (a) Micorscopic mages of lipid droplets in pre-adipocytes (day 0) and differentiated mature adipocytes (day 12) visualised using microscopy (x10 magnification - scale bar 10 µM) (b) Expression of key adipose genes during a 12-day differentiation period (normalised to housekeeping gene B2M). For each gene, the fold-change compared to the day expression is first observed is presented. Data represent three biological replicates and plotted as the mean ± SEM.

### Slide 2
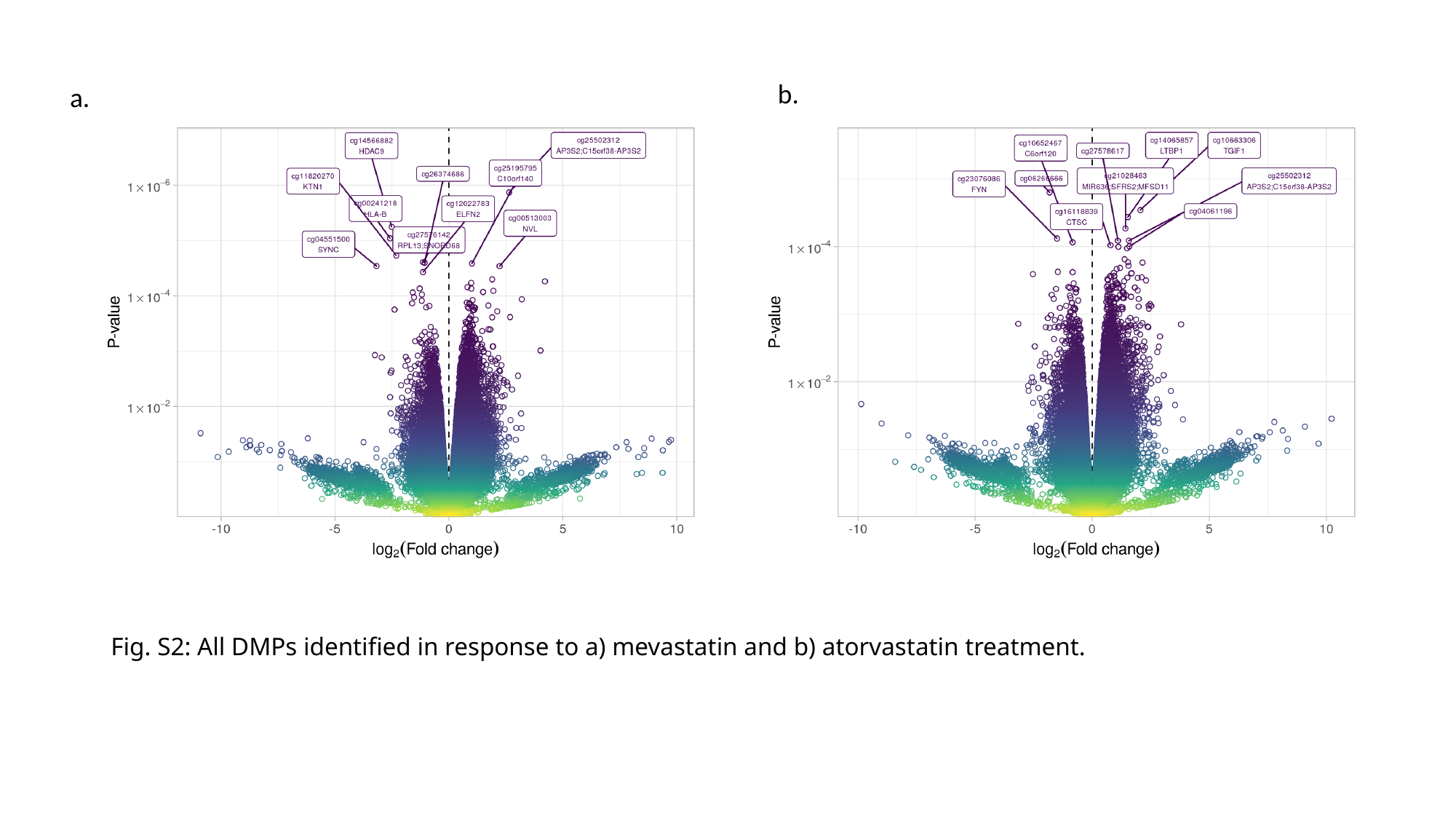

b.
a.
#### Fig. S2: All DMPs identified in response to a) mevastatin and b) atorvastatin treatment.

### Slide 3
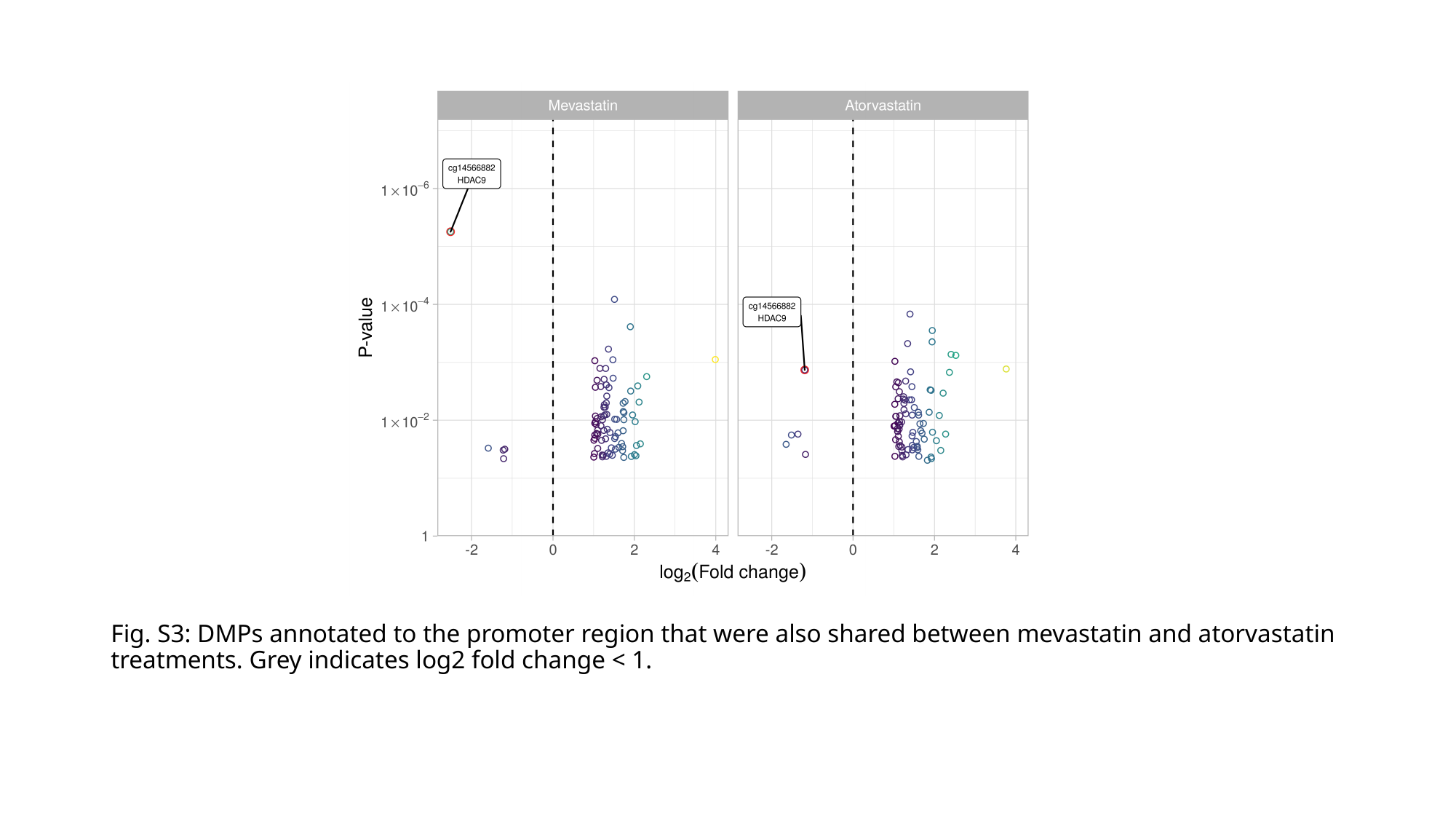

Fig. S3: DMPs annotated to the promoter region that were also shared between mevastatin and atorvastatin treatments. Grey indicates log2 fold change < 1.

### Slide 4
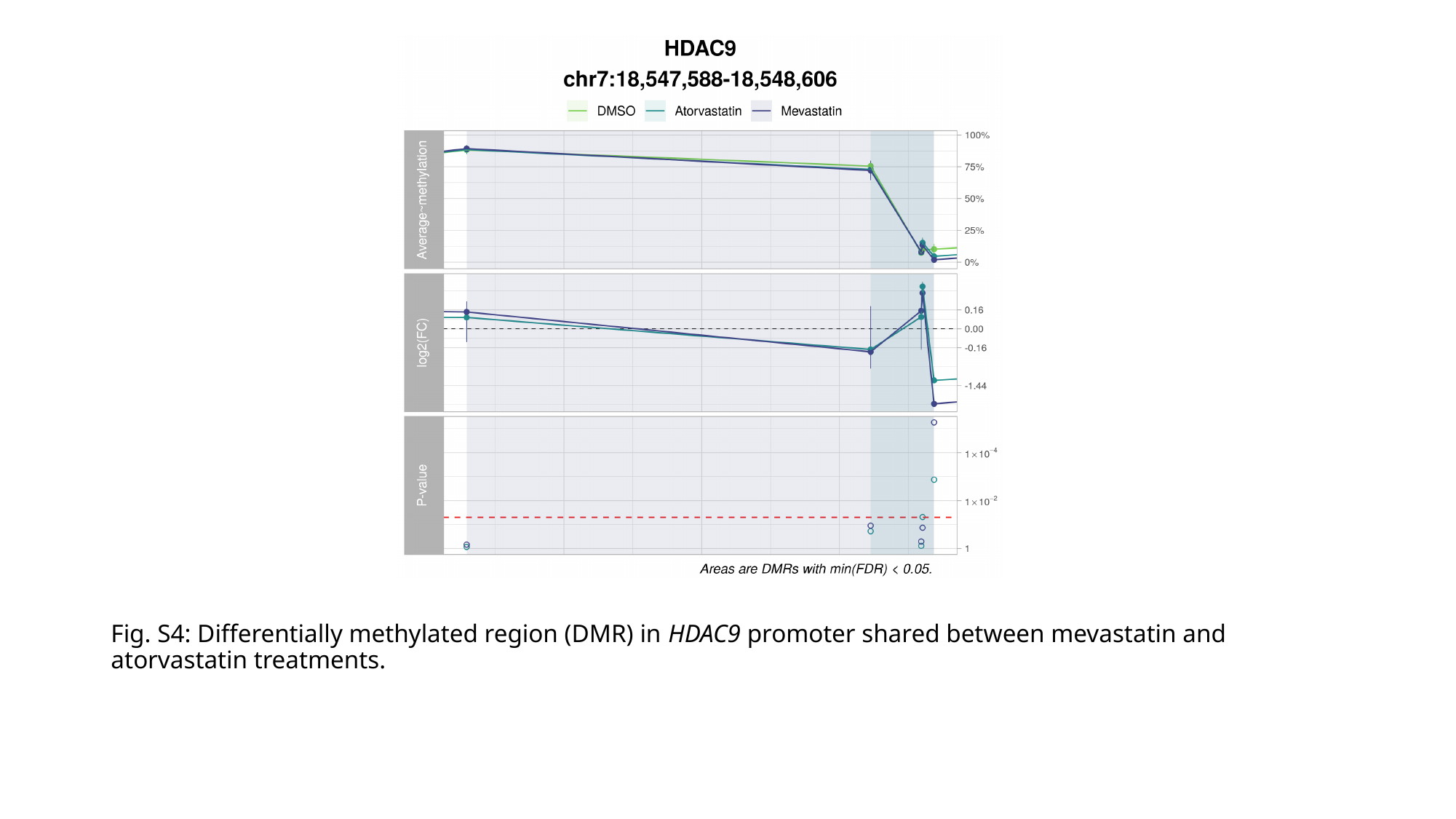

Fig. S4: Differentially methylated region (DMR) in HDAC9 promoter shared between mevastatin and atorvastatin treatments.
